# Common Pitfalls in CircRNA Detection and Quantification

**DOI:** 10.64898/2026.02.02.703185

**Authors:** Malte Weyrich, Nico Trummer, Fabian Böhm, Priscilla A. Furth, Markus Hoffmann, Markus List

**Affiliations:** Data Science in Systems Biology, School of Life Sciences, Technical University of Munich, Freising, Germany; Georgetown University, Washington, DC, USA; Department of Biochemistry and Molecular & Cellular Biology, Georgetown University Medical Center, Washington, DC, USA

**Keywords:** circRNA detection, RNA sequencing, bioinformatics, quantification, benchmarking

## Abstract

1

Circular RNAs have garnered considerable interest, as they have been implicated in numerous biological processes and diseases. Through their stability, they are often considered promising biomarker candidates or therapeutic targets. Due to the lack of a poly(A) tail, circRNAs are best detected in total RNA-seq data after depleting ribosomal RNA. However, we observe that the application of circRNA detection in the vastly more ubiquitous poly(A)-enriched RNA-seq data still occurs. In this study, we systematically compare the detection of circRNAs in two matched poly(A) and ribosomal RNA-depleted data sets. Our results indicate that the comparably few circRNAs detected in poly(A) data are likely false positives. In addition, we demonstrate that the quality of sample processing, as measured by the fraction of ribosomal reads, significantly affects the sensitivity of circRNA detection, leading to a bias in downstream analysis. Our findings establish best practices for circRNA research: total RNA sequencing with effective rRNA depletion is the preferred approach for accurate circRNA profiling, whereas poly(A)-enriched data are unsuitable for comprehensive detection. Employing multiple circRNA detection tools and prioritizing back-splice junctions identified by several algorithms enhances confidence in the selection of candidates. These recommendations, validated across diverse datasets and tissue types, provide generalizable principles for robust circRNA analysis.

**Key Points:**

- Ribosomal RNA contamination substantially impairs the accuracy of circRNA detection. This technical confounding factor has thus far received limited attention in the field.
- Tool agreement for circRNA calls is moderate in total RNA-seq but essentially absent in poly(A)-enriched RNA-seq data, underscoring the importance of using multiple tools for circRNA detection.
- Back-splice junctions detected in poly(A)-enriched RNA-seq data are predominantly tool-specific artifacts rather than genuine circRNAs, challenging the validity of circRNA identification in poly(A)-enriched datasets.

## 2 Introduction

Circular RNAs (circRNAs) are a class of RNA molecules characterized by a covalently closed loop structure that results from a non-canonical splicing event known as back-splicing, in which a downstream 5’ splice site is joined to an upstream 3’ splice site [14]. This circular configuration renders circRNAs resistant to exonuclease-mediated degradation, distinguishing them from linear RNAs. CircRNAs play important regulatory roles in gene expression, notably through microRNA sponging [12, 4] and by participating in the competing endogenous RNA network alongside other RNA species [11]. Dysregulation of circRNAs has been implicated in a wide range of diseases, including various cancers and neurological disorders [26].

Detection of circRNAs typically relies on identifying reads spanning back-splice junctions (BSJs) in RNA sequencing data [20]. However, because back-splicing is inherently less efficient than canonical linear splicing [9], circRNAs are usually expressed at lower levels. Moreover, only few reads originate from BSJs, making the distinction between the ubiquitous linear and the lowly expressed circRNAs computationally challenging. CircRNA detection is commonly applied to total RNA-seq libraries, as the circular structure of circRNAs lacks the poly(A) tail characteristic of most linear mRNAs. Notably, circRNAs are thus theoretically absent from poly(A)-selected RNA-seq datasets. In total RNA-seq protocols, it is common to employ ribosomal RNA (rRNA) depletion, as rRNA is the most abundant RNA type in the cell. CircRNAs can also be specifically enriched by RNase R digestion of linear transcripts to further support their detection [28][22].

While circRNAs are expected to be depleted in poly(A) libraries due to their lack of poly(A) tails, some circRNAs remain detectable in these libraries, either due to incomplete selection or the occurrence of A-rich sequences [23]. Analyses of non-colinear transcripts have shown that circRNAs can be detected in both poly(A)- and nonpoly(A)-selected data, though distinguishing them from other transcript types remains challenging [6]. Recent large-scale tool comparisons have been conducted primarily using total RNA or rRNA-depleted data [29, 27].

Although circRNAs are not necessarily expected to occur in poly(A)-enriched data sets, it is relatively common to apply computational identification of circRNAs, likely due to the widespread availability of large-scale poly(A)-selected transcriptomic resources such as the Cancer Cell Line Encyclopedia (CCLE) and The Cancer Genome Atlas (TCGA) [19]. The practical appeal of leveraging these extensive repositories for circRNA discovery is significant, as generating matched total RNA-seq datasets would be resource-intensive. However, this strategy is problematic. For instance, *Ruan et al*. [21] detected circRNAs in poly(A)-selected data using multiple computational tools and a consensus-based approach, yet acknowledged that the overlap between circRNAs detected in total RNA and poly(A)-enriched samples is typically small, both across datasets and between detection tools, raising concerns about false positive rates. Similarly, *Van der Steen et al*. [25] performed a related analysis using only a single detection tool (CIRCexplorer2), which limits the confidence in circRNA calls derived from poly(A)-selected data compared to using multiple algorithms.

The aim of our study is two-fold: (i) we perform the first systematic comparison of circRNA detection in matched poly(A)-enriched versus total RNA-seq data across different tools to assess the reliability and the practical utility of identifying circRNAs in poly(A)-selected datasets; (ii) we investigate if sample processing, i.e. rRNA depletion or poly(A) enrichment, introduces a sample-specific bias that leads to false positives in differential circRNA detection between sample groups. In this study we systematically assess these issues, describe pitfalls of circRNA detection and quantification and offer guidance on best practices.

The overall strategy of our study is illustrated in Figure 1, highlighting the key components and their relationships.

**Figure 1.**
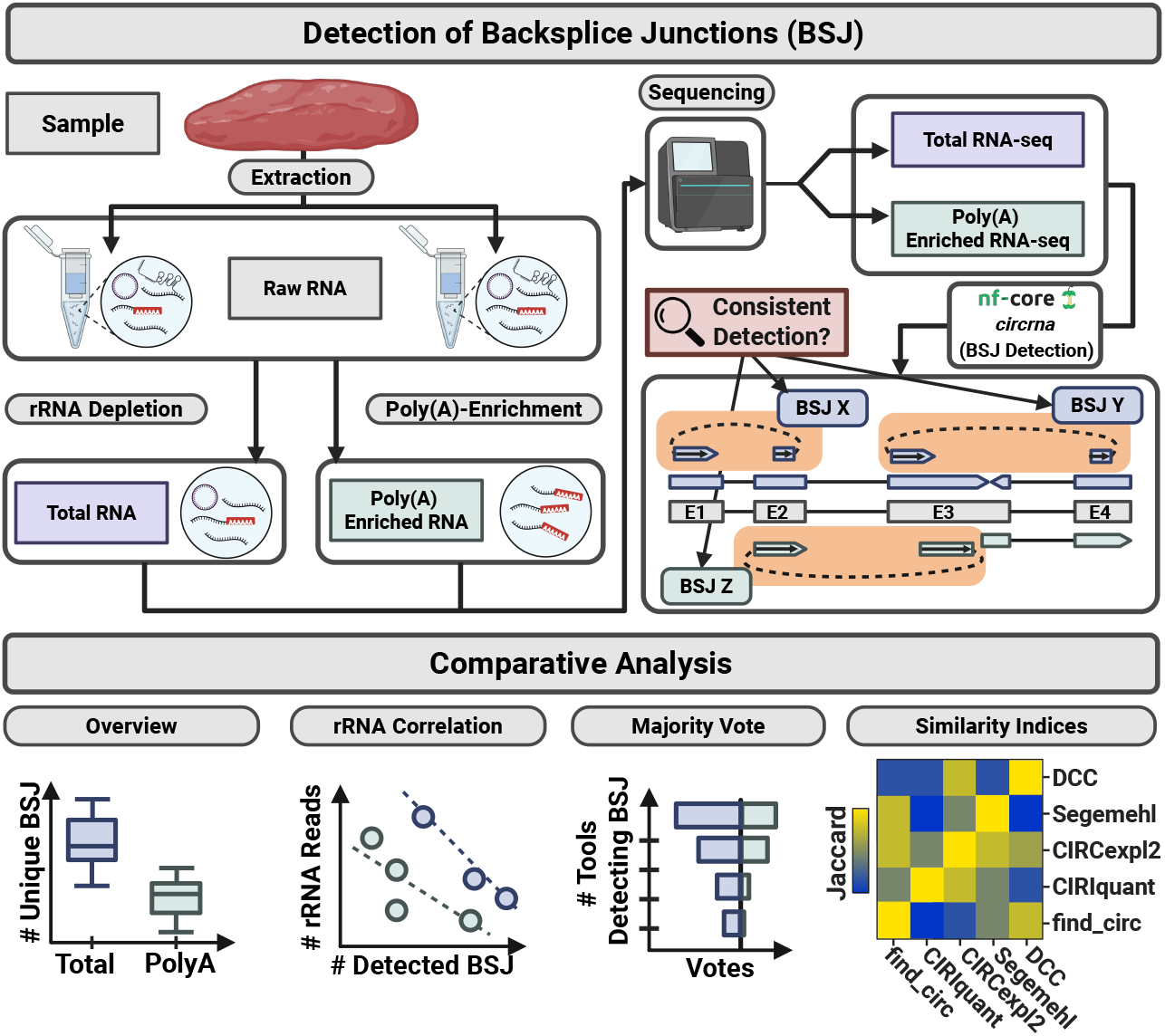
Overview of circRNA detection and comparative analysis workflow. Samples are processed for RNA extraction, yielding raw RNA. Two library preparation strategies are applied: total RNA after rRNA depletion (left) and Poly(A)-enriched RNA (right), producing respective RNA-seq libraries. Libraries are sequenced, generating total RNA-seq and Poly(A)-enriched RNA-seq data. Backsplice junctions (BSJs) are identified using the `nf-core/circrna` pipeline, producing BSJ calls for each library. Consistency across sequencing strategies and pipelines is assessed. The lower panel summarizes comparative analyses: (1) number of unique BSJs detected in total versus Poly(A) RNA; (2) correlation between rRNA reads and detected BSJs; (3) majority voting among multiple circRNA detection tools; and (4) similarity indices (e.g. Jaccard) comparing BSJ calls across tools (DCC, Segemehl, CIRCexpl2, CIRIquant, find circ). This illustration was created using BioRender [3].

## 3 Methods

### 3.1 CircRNA Detection

CircRNA detection was performed using the `nf-core/circrna` (v0.0.1dev) [8, 12] pipeline across two different datasets that provided both total and poly(A)-enriched RNA-seq (see Section 3.4 [24]): DEEP poly(A)-enriched vs total RNA [7], as well as GSE138734 poly(A)-enriched vs GSE138734 [17] total RNA datasets. This standardized workflow enabled consistent processing and comparison across different RNA preparation methods and datasets.

An advantage of the `nf-core/circrna` pipeline is that it employs five different circRNA detection tools to maximize sensitivity and confidence in BSJ identification: CIRI-quant [30], CIRCexplorer2 [31], find circ [18], DCC [5], and Segemehl [13]. Each tool implements distinct algorithmic approaches for identifying back-splice junctions, and their combined use allows for cross-validation of circRNA candidates. The pipeline processed each sample independently, generating BAM files and tool-specific BED files containing detected BSJ coordinates (see Figure 2.1). These standardized output files were extracted from the pipeline and served as the foundation for all subsequent filtering, merging, and comparative analyses described in the following sections.

**Figure 2.**
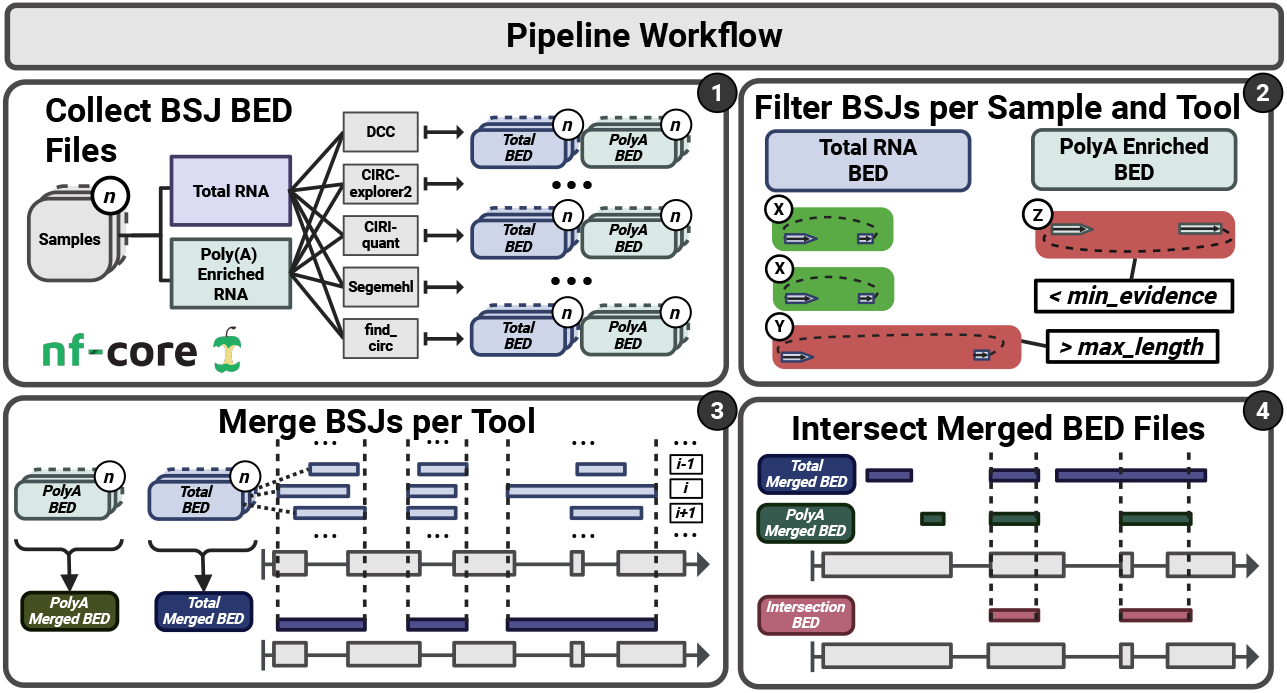
Pipeline workflow for BSJ detection using the `nf-core/circrna` [8] pipeline (1), followed by post-processing steps including: (2) removal of low-evidence or extremely long BSJs, (3) merging of BED files per tool, and (4) intersection of results across tools. This illustration was created using BioRender [3].

During the execution of the pipeline, the default parameters were used together with a blacklist [1] to prevent BSJ artifacts caused by known repeats in the human genome.

### 3.2 Post Processing

#### 3.2.1 Filtering

Each resulting BED file was filtered based on two quality criteria to remove likely artifacts and low-confidence calls. First, BSJs spanning more than 100,000 bp were excluded, as extremely long circRNAs are rare and likely represent false positives or mapping errors. Second, BSJs supported by fewer than 5 reads were removed to ensure only well-supported candidates were retained for downstream analysis, a threshold at which circRNA detection tools show improved precision [27].

#### 3.2.2 Merging and Intersection

Following filtering, BED files were systematically merged to identify unique BSJs across samples and tools. Initially, BED files from all samples within each dataset and RNA type were merged on a per-tool basis, yielding five BED files per dataset and type combination (one for each detection tool). Each of these files contained the combined set of unique BSJs detected by that tool across all samples. These tool-specific BED files were then merged again to obtain the total number of unique BSJs detected per dataset and RNA type, resulting in four comprehensive BED files (two per dataset: one for poly(A)-enriched and one for total RNA).

To assess the concordance between detection tools and RNA preparation methods, the tool-specific BED files were intersected in pairwise comparisons within each dataset. These comparisons included: (1) total RNA versus total RNA BSJs across different tools, (2) poly(A)-enriched versus poly(A)-enriched BSJs across different tools, and (3) total RNA versus poly(A)-enriched BSJs to evaluate the effect of the library preparation method on circRNA detection.

#### 3.2.3 rRNA Quantification

To assess ribosomal RNA (rRNA) contamination, we quantified rRNA-spanning reads using featureCounts (v2.0.6) [15] with a comprehensive rRNA annotation GTF file [32]. Reads were counted in paired-end mode (`-p --countReadPairs`), with multi-mapping reads included and fractionally assigned (`-M --fraction`). Counting was performed at the exon level (`-t exon`) and summarized by gene identifier (`-g gene _id`). Read counts were normalized to counts per million (CPM) for each sample across both datasets and RNA types to enable direct comparison of rRNA content between samples.

### 3.3 Comparison of Detected CircRNAs in Poly (A) and Total RNA-seq

#### 3.3.1 Detection Concordance Analysis

To evaluate the consistency of circRNA detection across different library preparation methods and bioinformatic tools, we performed multi-level comparisons of unique BSJs. First, we compared the total number of unique BSJs between poly(A)-enriched and total RNA datasets using the merged per-dataset BED files. Second, we assessed tool-level concordance by determining how many BSJs were detected by multiple tools within each dataset and RNA type. For each BSJ, we calculated a support score ranging from 1 to 5, where a score of 5 indicates detection by all five tools (representing maximum consensus), while lower scores reflect detection by fewer tools. This majority vote approach enabled us to stratify circRNA candidates by detection confidence.

#### 3.3.2 rRNA Contamination Analysis

To assess the potential impact of rRNA contamination on circRNA detection, we examined the relationship between rRNA content and BSJ identification. For each tool, the total number of BSJ-spanning reads was compared to the number of rRNA-spanning reads in the same sample. Pearson correlation coefficients were calculated per tool and RNA type to quantify the strength and direction of any association between rRNA contamination and apparent circRNA detection levels.

#### 3.3.3 Tool Similarity Metrics

To quantify the similarity and overlap between different detection tools, we employed two complementary metrics on the per-tool BED files. First, we calculated the Jaccard index to measure overall similarity between tool outputs using `bedtools jaccard:`

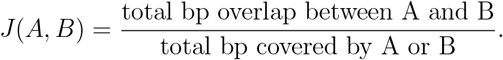

Second, we computed intersection percentages to capture asymmetric detection patterns. The intersection percentage represents the proportion of BSJs from one tool that overlap with BSJs detected by another tool, normalized by the total number of unique genomic loci detected by both tools:

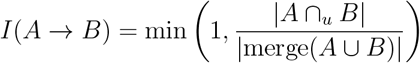

where |*A* ∩_*u*_ *B*| denotes the count of intervals from tool *A* that overlap with at least one interval from tool *B* (obtained via `bedtools intersect -u -a A -b B`), and |merge(*A* ∪ *B*) | represents the number of unique, non-overlapping genomic regions obtained by concatenating both BED files, sorting, and applying `bedtools merge`. Due to coordinate variations in how different tools report the same BSJ, the merge operation occasionally resulted in fewer unique regions than intersecting intervals; in such cases, the metric was capped at 1.0 to reflect complete overlap. This metric is asymmetric, as the proportion of *A* overlapping with *B* may differ from *B* overlapping with *A*, thereby revealing directional detection patterns and tool-specific sensitivities. Pairwise comparisons across all tools and samples are shown in Supplementary Figure 5.

### 3.4 Code and Data Availability

The RNA-seq datasets analyzed in this study are publicly available.

#### 3.4.1 GSE138734 Dataset

Paired-end RNA-seq data derived from neuroblastoma (sympathetic nervous system) cell lines, including 10 manually selected pairs of poly(A)-enriched and total RNA samples, were downloaded from the Gene Expression Omnibus (GEO [2], accession GSE138734 [10]) using the SRA Toolkit command:

`fastq-dump <SRR-ID> --split-files -O [polya|total] fastq/`. This dataset is described in the related publication by [16].

#### 3.4.2 DEEP Dataset

RNA-seq data was obtained from the DEEP project (http://www.deutsches-epigenom-programm.de/). using the the dataset EGAD00001002735 [7] hosted by the European Genome-Phenome Archive (https://ega-archive.org/). A total of thirteen paired-end FASTQ files were manually selected for each RNA category (poly(A) and total RNA for each sample), comprising: four hepatocyte (LiHe), three naive CD4-positive T cell (BlTN), three effector memory CD4-positive alpha-beta T cell (BlEM), and three central memory CD4-positive alpha-beta T cell (BlCM) samples.

#### 3.4.3 Code of Analysis

All relevant scripts can be found at https://github.com/daisybio/circRNAbenchmarking.

## 4 Results and Discussion

The comparison of circRNA detection between total RNA-seq and poly(A)-enriched data across two datasets (DEEP and GSE138734) revealed substantial differences in detection efficiency and tool performance (see Figure 3). Total RNA-seq consistently identified significantly more unique BSJs than poly(A)-enriched samples: 4,800 versus 384 BSJs in DEEP and 2,161 versus 317 in GSE138734, representing 12.5-fold and 7-fold differences, respectively (Sub-plot 3(a)). Notably, despite DEEP having approximately double the SAM entries compared to GSE138734 (780 million versus 347 million for total RNA-seq), BSJ detection in total RNA-seq also increased by two-fold, while poly(A) data showed only minimal gains with increased sequencing depth, indicating that poly(A)-enriched libraries have an inherently weaker biological signal for circRNAs. Across both datasets, Segemehl detected the highest number of unique BSJs for both data types, while DCC, CIRIquant, and find circ showed similar detection levels in DEEP but varying amounts in GSE138734 (Sub-plot 3(b)(d)). All tools struggled with poly(A)-enriched data, detecting very few circRNAs regardless of their algorithm.

**Figure 3.**
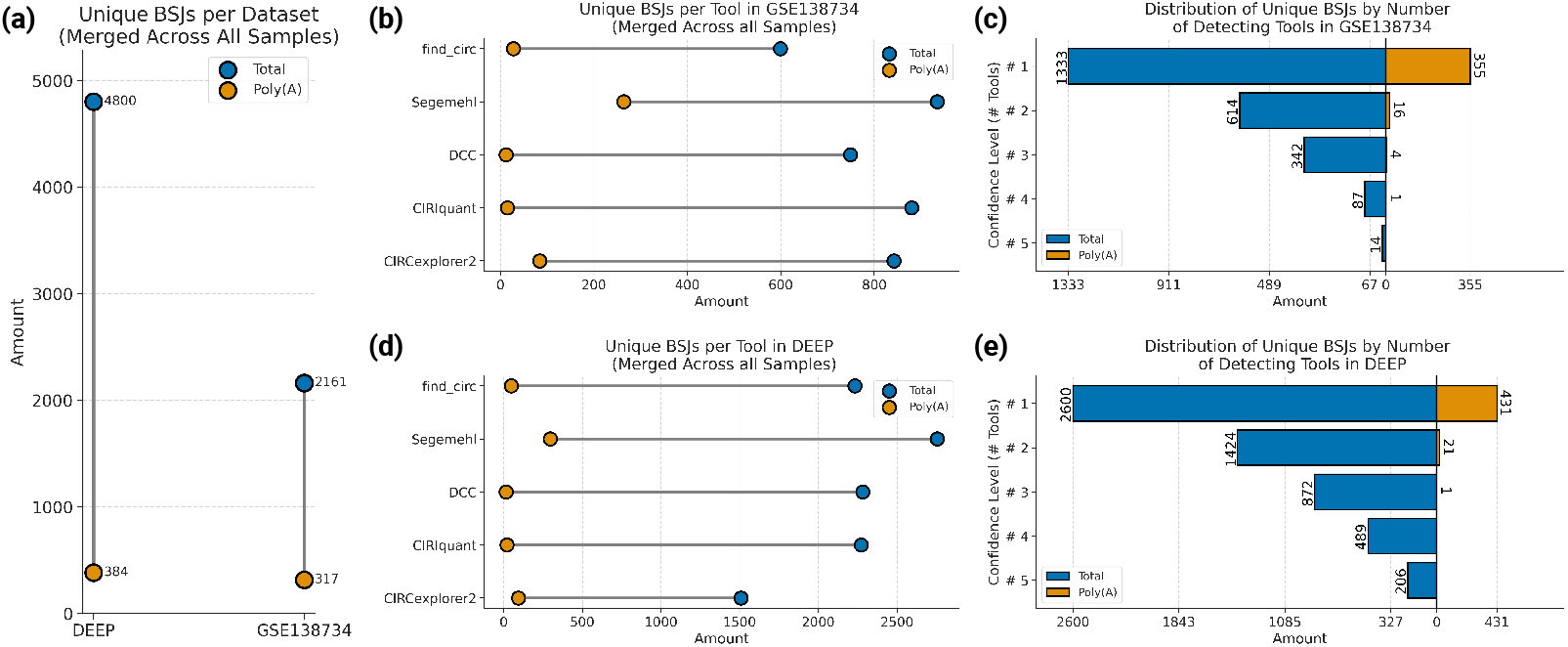
Comparison of circRNA detection between total RNA-seq and poly(A)-enriched data across DEEP and GSE138734 datasets. (a) Total BSJs per dataset merged across samples and tools. (b, d) Tool-specific detection in GSE138734 and DEEP, respectively. (c, e) Distribution of BSJs by number of detecting tools (confidence levels 1-5) in GSE138734 and DEEP, respectively. Total RNA-seq consistently shows higher detection rates and better inter-tool agreement compared to poly(A)-enriched data.

An analysis of detection confidence levels revealed important patterns in tool agreement and reproducibility (Figure 3(c)(e)). The majority of BSJs were detected by only a single tool (confidence level 1), with subsequent confidence levels showing progressively fewer BSJs as tool agreement increased. For total RNA-seq, the number of BSJs roughly halved at each increasing confidence level, suggesting moderate overlap between tools. However, poly(A)-enriched data showed drastically reduced tool agreement beyond confidence level 1, indicating that the limited circRNAs detected in poly(A) libraries are highly tool-specific. Only a small fraction of BSJs were detected by two tools (confidence level 2): 16 in GSE138734 and 21 in DEEP. Notably, almost no BSJs in poly(A)-enriched data were detected by more than two tools, indicating minimal multi-tool agreement. These patterns were consistent across both datasets, and detailed tool intersections are provided in Supplementary Figure 3. These findings demonstrate that total RNA-seq is the preferred method for circRNA studies, as poly(A) enrichment, as expected, systematically depletes circRNAs and reduces detection reproducibility across tools.

In both datasets, total RNA-seq libraries showed significant negative correlations (see Figure 4) between rRNA contamination and BSJ detection (except for find circ in the DEEP dataset), indicating that rRNA contamination reduces the remaining sequencing depth for circRNA reads. This relationship is observed in both datasets, which have varying amounts of rRNA-spanning reads (see Supplement Figure 1). While DEEP contains substantially more rRNA reads than GSE138734 in total RNA-seq, both show negative correlations because the higher volume of available circRNAs in total RNA-seq makes BSJ detection sensitive to even modest rRNA contamination.

**Figure 4.**
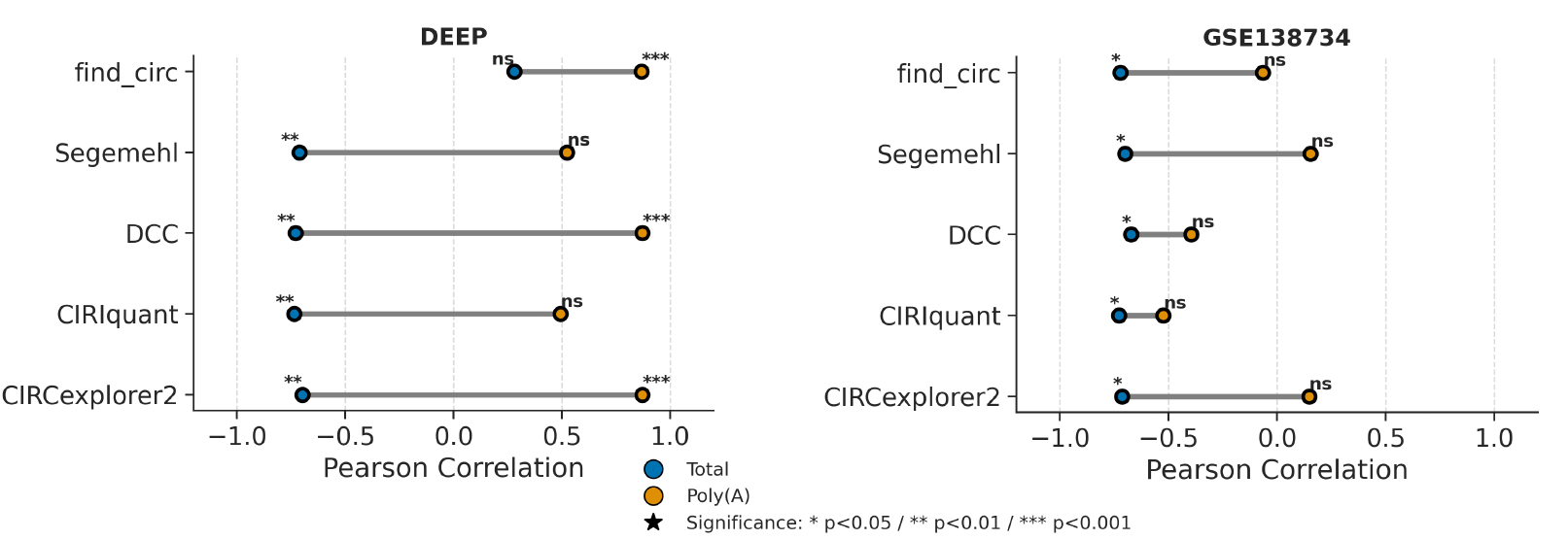
Impact of rRNA contamination on circRNA detection across sequencing methods and datasets. Dumbbell plots showing Pearson correlation coefficients between rRNA-spanning read counts and sum of BSJ supporting reads for each tool in DEEP (left) and GSE138734 (right) datasets. In total RNA-seq, negative correlations indicate that in-creased rRNA contamination reduces BSJ detection, with stronger effects in non-rRNA-depleted samples (DEEP). Poly(A)-enriched samples show mixed and mostly insignificant correlations, reflecting naturally lower rRNA content. The apparent positive correlation in DEEP poly(A) samples is likely a technical artifact (see Supplementary Figure 1 and 2 for detailed analysis).

In contrast, poly(A)-enriched samples showed mixed and non-significant correlations between rRNA reads and BSJ detection across both datasets. While rRNA depletion occurs naturally during poly(A) selection as non-polyadenylated transcripts are excluded, residual rRNA contamination can still occur due to incomplete enrichment or cross-contamination. The apparent positive correlation observed in DEEP poly(A) samples is likely a technical artifact arising from multiple confounding factors which are further investigated in Supplementary Figures 1 and 2.

These findings demonstrate that rRNA contamination in total RNA-seq libraries negatively impacts circRNA detection by competing for sequencing depth, emphasizing the importance of effective rRNA depletion for circRNA studies.

To assess tool concordance, we calculated Jaccard indices comparing the merged BED files for each tool across both datasets and sequencing methods. In total RNA-seq, the Jaccard index was frequently high for most tool pairs in both datasets (see Figure 5 (a)), with DCC and CIRCexplorer2 showing the highest similarity across both GSE138734 and DEEP. Segemehl repeatedly showed lower similarity to other tools due to detecting substantially more BSJs, resulting in larger union sets and thus lower Jaccard indices. In poly(A)-enriched data, tool similarity patterns differed between datasets (see Figure 5 (b)): GSE138734 maintained high Jaccard indices among CIRIquant, CIRCexplorer2, and DCC, while DEEP showed reduced similarity across all tools except for Segemehl, which remained the most dissimilar. However, examination of BSJ length distributions revealed that these high Jaccard indices can be misleading, as they are driven by longer BSJs that dominate total genomic coverage, while many shorter BSJs remain tool-specific and do not overlap (Supplementary Figure 4). Cross-datatype comparisons between poly(A) and total RNA-seq (see Figure 5 (c)) showed consistently low Jaccard indices across all tools in both datasets, reflecting the fundamental differences in circRNA detection between these methods. While GSE138734 exhibited slightly higher cross-datatype similarity than DEEP, this difference is primarily attributable to the substantially higher volume of BSJ detections in DEEP, which increases the union set and consequently reduces the Jaccard index. These findings demonstrate that DCC and CIRCexplorer2 show the most consistent overlap across datasets and data types, while the Jaccard metric’s sensitivity to both BSJ length and detection volume must be considered when interpreting tool concordance.

**Figure 5.**
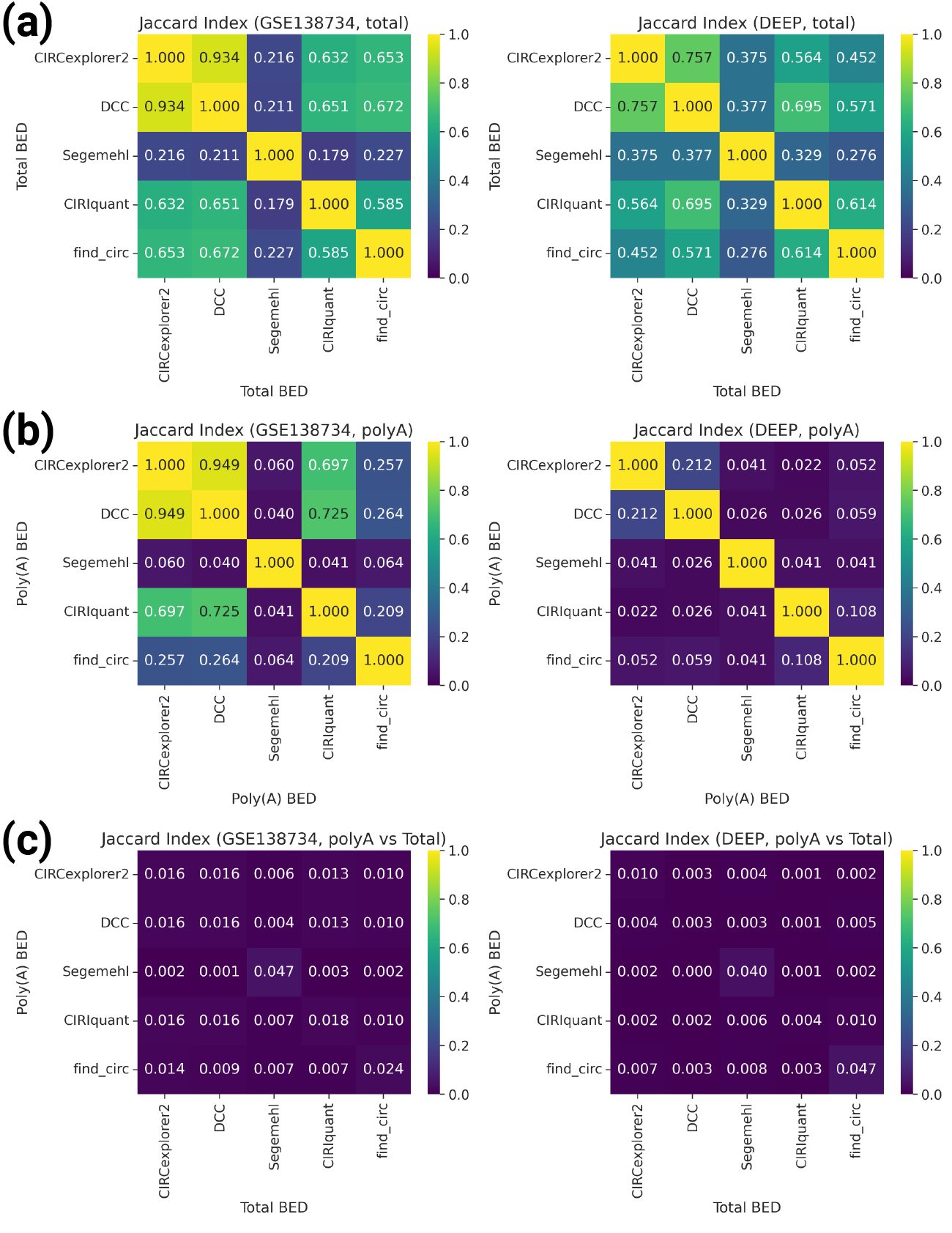
Tool concordance assessed by Jaccard index across datasets and sequencing methods. Heatmaps showing pairwise Jaccard indices between merged BED files for each detection tool. (a) Tool comparisons within total RNA-seq for GSE138734 (left) and DEEP (right), showing high similarity between DCC and CIRCexplorer2, with Segemehl consistently showing lower overlap due to detecting more BSJs. (b) Tool comparisons within poly(A)-enriched data for GSE138734 (left) and DEEP (right), showing dataset-specific patterns with GSE138734 maintaining high tool concordance while DEEP shows reduced similarity. (c) Cross-datatype comparisons between poly(A) and total RNA-seq for GSE138734 (left) and DEEP (right), demonstrating consistently low similarity between sequencing methods across all tools. Jaccard index values range from 0 (no overlap) to 1 (complete overlap).

## 5 Limitations and Considerations

Several technical and biological factors should be considered when interpreting the results of this study. First, the sample sizes are relatively modest, with 13 samples in DEEP and 10 samples in GSE138734, which may limit statistical power for detecting correlations and assessing significance. Second, the DEEP dataset comprises heterogeneous tissue types: 4 liver samples (LiHe), and 9 blood-derived samples from three T-cell subgroups (BlTN, BlEM, and BlCM); which introduces biological variability that could confound analyses comparing technical effects across samples.

The rRNA depletion efficiency also varies substantially between datasets. In DEEP, the total RNA-seq samples, which should have undergone rRNA depletion, paradoxically contain many more rRNA-spanning reads compared to poly(A)-enriched samples (Supplementary Figure 1), raising questions about the effectiveness or consistency of rRNA depletion protocols. In contrast, GSE138734 shows the expected pattern, with total RNA-seq samples (post-depletion) containing fewer rRNA reads than would be expected without depletion. These differences in rRNA content may influence the observed correlations between rRNA contamination and BSJ detection.

Additionally, tool-specific artifacts were observed in the DEEP dataset. Notably, find circ showed abnormally high BSJ evidence scores for certain samples—particularly the liver samples—with total supporting read counts that exceeded even those in total RNA-seq data (Supplementary Figure 2). This pattern was unique to find_circ and not observed with other tools, suggesting a tool-specific sensitivity or artifact that could influence correlation analyses and significance testing. These tool- and sample-specific effects highlight the importance of using multiple detection tools and carefully examining individual sample behavior when assessing circRNA detection patterns.

## 6 Conclusion

This comprehensive comparison of circRNA detection across two independent datasets (DEEP and GSE138734) and two sequencing methodologies (total RNA-seq and poly(A) enrichment) reveals fundamental differences in circRNA detectability and tool performance that have important implications for experimental design and data interpretation.

Our findings demonstrate that total RNA-seq is superior for circRNA discovery and quantification. Across both datasets, total RNA-seq detected 7- to 12.5-fold more unique back-splice junctions than poly(A)-enriched libraries, and this difference is not simply a matter of sequencing depth, doubling the number of sequenced reads in poly(A) data yielded minimal gains in circRNA detection. This poor scaling reflects the fundamental biology of circRNAs: most, if not all, lack poly(A) tails and are therefore systematically depleted during poly(A) selection. The few circRNAs detected in poly(A) libraries show substantially reduced inter-tool agreement and more tool-specific detection patterns, suggesting that many may represent technical artifacts rather than a robust biological signal. Consequently, we strongly advise against the practice of detecting circRNAs in poly(A)-enriched data, as the detected circRNAs (typically, studies employ only a single method) are likely to be false positives, rendering any conclusions drawn questionable.

Ribosomal RNA contamination emerged as a significant confounding factor in total RNA-seq libraries, with clear negative correlations between rRNA-spanning reads and BSJ detection across multiple tools in both datasets. This relationship persists even in libraries that underwent rRNA depletion, emphasizing that effective rRNA removal is crucial for maximizing the sensitivity of circRNA detection. The competition for sequencing depth between rRNA and circRNA-spanning reads is particularly impactful in total RNA-seq because of the higher overall abundance of circRNAs in these libraries. Consequently, ribosomal RNA contamination leads to sample-specific biases in the expression of circRNAs that are not countered by normalizing for sequencing depth. This bias likely affects differential expression analysis between sample groups if ribosomal RNA content between the groups differs systematically. To mitigate this bias, we advise including the fraction of ribosomal RNA as a confounder in differential expression analysis.

Tool selection and multi-tool validation are critical considerations for circRNA studies. While all five tools (Segemehl, DCC, CIRCexplorer2, CIRIquant, and find circ) showed consistent detection patterns across datasets, they exhibited substantial differences in sensitivity and specificity. Segemehl consistently detected the most BSJs, while DCC and CIRCexplorer2 showed the highest concordance in DEEP, suggesting they may identify a more conserved set of high-confidence circRNAs. Importantly, the majority of detected BSJs were identified by only a single tool, while just 14 (GSE138734)/206(DEEP) BSJs achieved the highest confidence level of detection by all five tools in total RNA-seq. This limited overlap underscores the importance of using multiple detection algorithms and establishing confidence thresholds based on multi-tool agreement.

The Jaccard index analysis revealed an important methodological feature: high genomic overlap (measured by base-pair coverage) can mask poor feature-level agreement when a small number of long BSJs dominate the overlap calculation. Our feature-level overlap analysis demonstrated that many shorter BSJs are tool-specific and do not overlap, despite high Jaccard indices suggesting strong concordance. This finding emphasizes the need for multiple complementary metrics when assessing tool performance and biological reproducibility.

From a practical standpoint, these results provide clear guidance for circRNA research: (1) total RNA-seq with effective rRNA depletion should be the method of choice for circRNA studies; (2) poly(A)-enriched data, while useful for mRNA analysis, should not be relied upon for comprehensive circRNA profiling; (3) multiple detection tools should be employed, with particular attention to BSJs detected by multiple algorithms as high-confidence candidates; and (4) rRNA contamination should be monitored and minimized or at least be accounted for in downstream analysis, as it directly impacts circRNA detection sensitivity. These recommendations are consistent across independent datasets and tissue types, suggesting they represent generalizable principles for circRNA detection and analysis.

## Biographical Note

M.W. is a scientific research student at the Technical University of Munich.

## Author Contributions

M.W. wrote the manuscript, developed the benchmarking code, and prepared the figures. F.B. contributed to the design of the benchmarking workflow. N.T. developed the `nf-core/circrna` pipeline and provided technical support. P.F., M.H., and M.L. provided guidance and supervision and reviewed the manuscript.

