## Supplementary Material for "Common Pitfalls in CircRNA Detection and Quantification"

### Supplement

Malte Weyrich[1], Nico Trummer[1], Fabian Böhm [1], Priscilla A. Furth[2]*, Markus Hoffmann[3]*, Markus List[1]*
(*joint last authors)

[1]*Data Science in Systems Biology, School of Life Sciences, Technical University of Munich, Freising, Germany*
[2]*Georgetown University, Washington, DC, USA*
[3]*Department of Biochemistry and Molecular & Cellular Biology, Georgetown University Medical Center, Washington, DC, USA*

**Corresponding author:**
Markus List
Data Science in Systems Biology
Technical University of Munich


**Funding:**
Funded by the Federal Ministry of Education and Research (BMBF) and the Free State of Bavaria under the Excellence Strategy of the Federal Government and the Länder, as well as by the Technical University of Munich – Institute for Advanced Study, Garching, Germany through an Anna Boyksen Fellowship (P.A.F.)

### Supplementary Figures

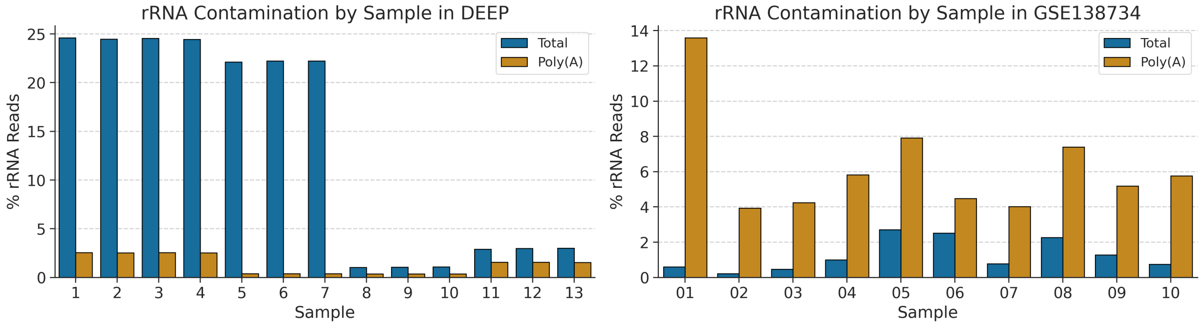


Figure 1: Comparison of share of rRNA spanning reads per dataset and RNA-seq type. The DEEP dataset has seven samples with abnormally high rRNA read ratios while samples eight to thirteen have a more reasonable volume. The opposite is true for GSE138734 where all poly(A)-enriched samples show a higher ratio compared to total RNA-seq. The apparent positive correlation between rRNA contamination and BSJ detection observed in DEEP poly(A) samples (main Figure 4) likely represents a technical artifact rather than a biological signal. Multiple confounding factors contribute to this pattern: (1) heterogeneous sample composition (four liver and eight T-cell samples from three groups with different biological characteristics: Samples 1–4: LiHe; 5–7: BlCM; 8–10: BlEM; 11–13: BlTN), (2) very low contamination of rRNA-spanning reads and a low number of BSJ detections in poly(A) libraries, making correlations unstable, (3) tool-specific artifacts, particularly abnormally high find_circ BSJ evidence scores in liver samples (see Supplementary Figure 2), and (4) differences in sequencing depth among sample subgroups. These factors collectively confound the relationship between rRNA content and circRNA detection in this specific subset of samples.

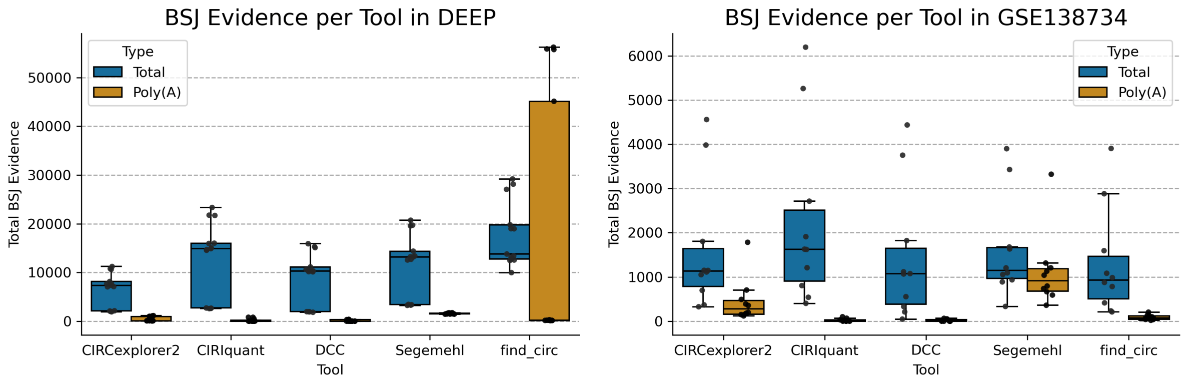


Figure 2: Total amount of BSJ supporting reads per tool and sample in both datasets. For each sample, the amount of BSJ supporting reads annotated in the BED file were summed up and compared between data types. Since DEEP has higher coverage compared to GSE138734, the evidence sum per sample is higher in all cases. The over all pattern of tools seems to stay consistent across datasets with the exception of find_circ, which shows abnormally high BSJ evidence specifically in DEEP samples (particularly samples 1–7, corresponding to liver samples with high rRNA contamination, see Supplementary Figure 1). This pattern is not observed in GSE138734, suggesting a dataset- or tissue-specific artifact.

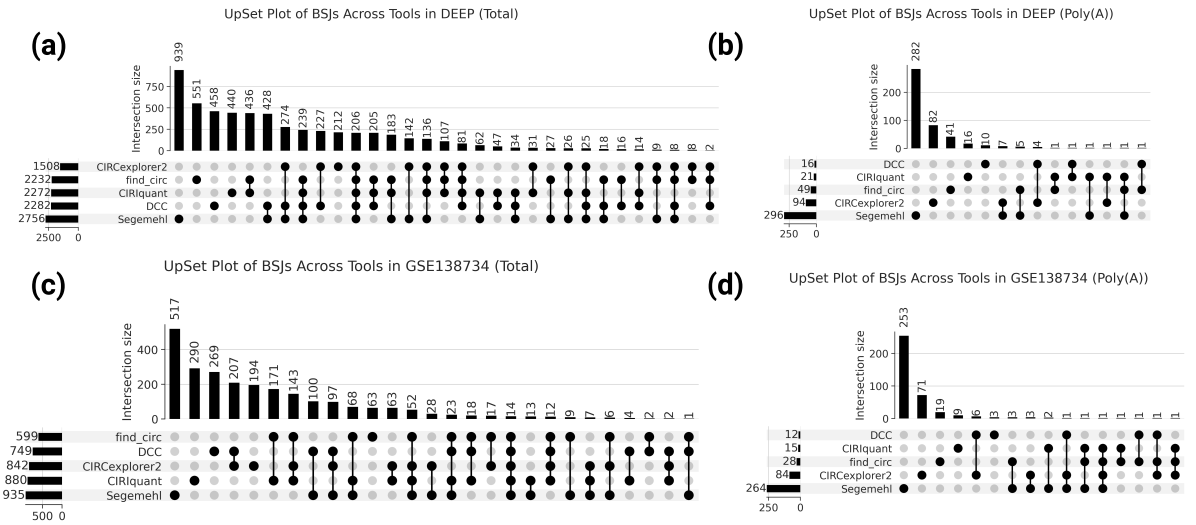


Figure 3: Upset plots of shared BSJs based on (chr, start, stop) across all tools in both datasets and data types. While there are many instances where several tools agree in total RNA-seq for both datasets, this cannot be said for poly(A)-enriched data. In DEEP there are 206 BSJs detected by all tools and 14 for GSE138734, yet there was not a single comparable BSJ in poly(A) data.

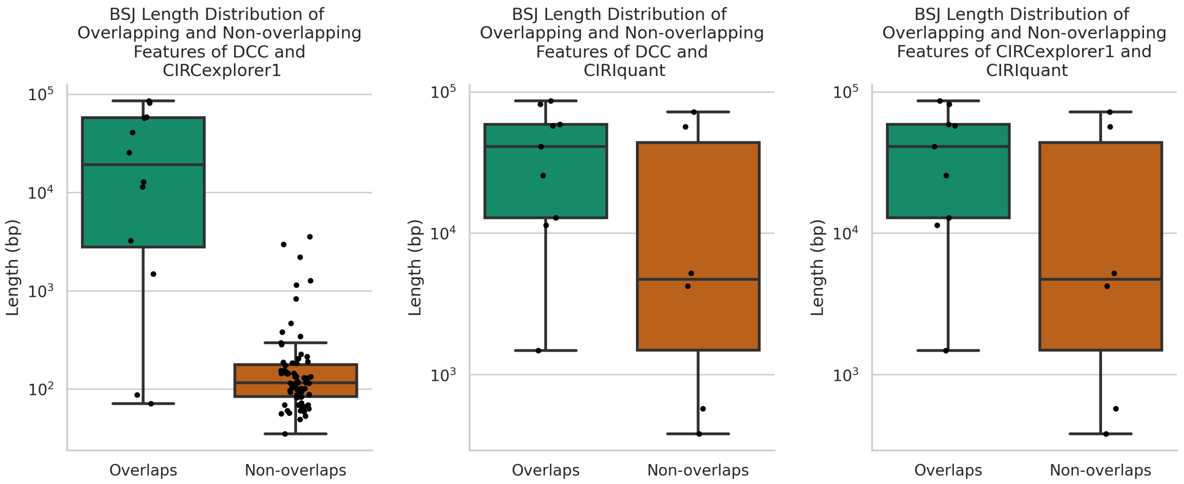


Figure 4: Length distribution of overlapping and non-overlapping BSJs. Since the Jaccard index seemed fairly high (0.697-0.949) for some poly(A)-enriched samples of the GSE138734 dataset (see Figure 5 in main paper), the lengths of the BSJs contributing to these scores were analyzed. Most notably was the Jaccard index of 0.949 between CIRCexplorer2 and DCC. This high similarity arises because the Jaccard index is strongly influenced by the lengths of overlapping intervals: long overlaps contribute disproportionately, while short, non-overlapping BSJs have little effect on the score. Even though the number of non-overlapping BSJs is greater, the long overlapping intervals dominate the Jaccard calculation. The same pattern is seen for CIRIquant vs. DCC and CIRCexplorer2 vs. CIRIquant, although to a lesser extent.

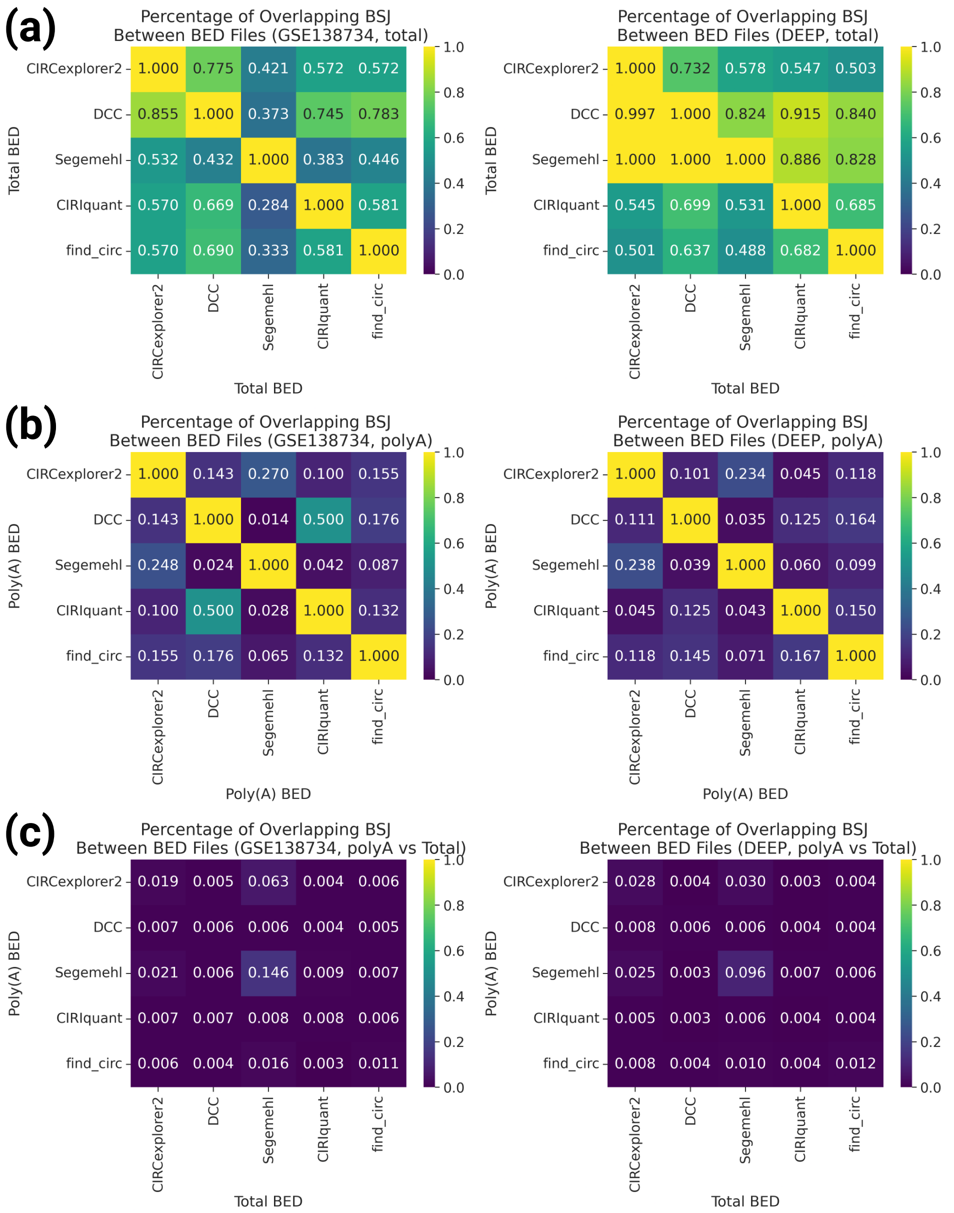


Figure 5: Similarity heatmaps showing the asymmetric relationship between certain tool pairs, revealing tool-specific detection patterns masked by the Jaccard index. To complement the Jaccard index analysis and address potential biases introduced by long BSJs dominating genomic coverage, we calculated the percentage of overlapping BSJs at the feature level rather than base-pair level.

In total RNA-seq, BSJ overlap percentages remained relatively high across both datasets (Figure 5 (a)), confirming that tools detect largely consistent sets of circRNAs, with DCC and CIRCexplorer2 showing the strongest concordance. However, in poly(A)-enriched data (5 (b)), the feature-level overlap was substantially lower than the Jaccard indices suggested, revealing that tools detect largely non-overlapping sets of BSJs. Across both datasets, only DCC and CIRIquant showed moderate overlap in poly(A) data, while other tool pairs exhibited minimal BSJ-level agreement, with overlap percentages often below 20%. The inter-datatype comparison between total RNA-seq and poly(A)-enriched RNA-seq in Figure 5(c) shows little to no overlap across the tools.

This striking contrast between high Jaccard indices and low feature-level overlap in poly(A) data confirms that the Jaccard metric was inflated by a small number of long, shared BSJs that dominate genomic coverage, while the majority of detected BSJs–particularly shorter ones–are tool-specific. Cross-datatype comparisons between poly(A) and total RNA-seq showed consistently minimal BSJ overlap (typically $< 5\%$) across all tools in both datasets, reinforcing the fundamental differences in circRNA detection between library preparation methods.

These findings suggest that the inconsistent BSJ detection patterns in poly(A)-enriched data, combined with minimal inter-tool agreement, may indicate that many poly(A)-detected regions represent tool-specific artifacts rather than robust biological signal. The feature-level overlap metric should thus always be considered together with the Jaccard index to identify cases where genomic coverage-based metrics may be biased by a small number of long BSJs.
